# Adaptive experiments in high-dimensional feature spaces: A particle filtering approach

**DOI:** 10.64898/2026.08.03.741989

**Authors:** Rabea Turon, Lars C. Reining, Philipp A. Hummel, Lynn Schmittwilken, Christine Lind, Angela J. Yu, Constantin A. Rothkopf, Frank Jäkel, Thomas S. A. Wallis

## Abstract

Behavioral experiments are often infeasible when stimulus spaces have many dimensions or when testing time is limited. One way to address this challenge is adaptive stimulus selection, where informative stimuli are chosen dynamically based on participants’ responses. However, in high-dimensional spaces, identifying such stimuli is computationally demanding. Here, we describe High-dimensional Online Particle Estimation (HOPE), which selects informative stimuli in less than a second for up to 50 dimensions, enabling efficient estimation of high-dimensional psychometric functions. We validate HOPE through simulations and a face-categorization experiment in an 18-dimensional parameter space with human participants. Compared to uniform stimulus presentation, HOPE reduces uncertainty over model parameters two-to three-times faster, reaching the same certainty in half the trials or fewer. This efficiency enables psychophysical studies that were previously impractical due to the exponential scaling of trial requirements.

## 1 Introduction

People can classify complex stimuli into categories, yet the mechanisms by which they do so remain poorly understood. In part, this is because complex stimuli can vary over many feature dimensions simultaneously (such as size, color, contrast, intensity, space or time). For example, to judge whether they perceive a face as male or female, humans mostly rely on the eye-eyebrow region and its luminance difference to other parts of the face (Dupuis-Roy et al., 2009, 2019; Macke and Wichmann, 2010; Russell, 2003). Psychometric functions quantify how decisions change as feature dimensions vary, yet standard methods for estimating psychometric functions become impractical as the number of dimensions grows. Selecting a fixed set of feature combinations to test, for instance, means that the required number of trials grows exponentially with the number of feature dimensions, translating into days or even weeks of data collection. This is often infeasible, particularly in clinical or other time-limited settings, necessitating methods that are both data-efficient and computationally tractable in high-dimensional spaces.

To improve efficiency, one can use efficient task designs (Jäkel and Wichmann, 2006; Bex and Sker-swetat, 2021; Neupane et al., 2024; Hou et al., 2015; Bonnen et al., 2015; Straub and Rothkopf, 2022; Skerswetat et al., 2024), and / or select stimuli adaptively based on participants’ responses, to concentrate data collection in the most informative regions of the stimulus space (Cornsweet, 1962; Watson and Pelli, 1983; King-Smith et al., 1994; Kontsevich and Tyler, 1999; Watson, 2017; Vul et al., 2010; Greenhill et al., 2020; Chaloner and Verdinelli, 1995; Foster et al., 2021; Barthelmé and Mamassian, 2008; DiMattina, 2015a; Lesmes et al., 2006, 2010; Park and Pillow, 2012; Prins, 2013; Lesmes et al., 2015; Kujala and Lukka, 2006; Bak and Pillow, 2018). Mostly, these methods select the next trial to minimize uncertainty over predefined model parameters. Because this is computationally intractable to calculate over continuous feature and parameter dimensions, most algorithms discretize the parameter space on a grid and use a fixed stimulus pool. This grid approximation normally grows exponentially with the stimulus feature dimensions. As a result, even highly efficient algorithms such as QUEST+ (Watson, 2017) become impractical beyond roughly five parameters.

Here, we further develop an adaptive sampling framework proposed by Kujala and Lukka (2006)^1^: we extend the framework to a large number of dimensions, optimize the algorithm so that it can be computed quickly, and empirically validate its performance both in simulation and in a real experiment. Our implementation, High-dimensional Online Particle Estimation (HOPE), integrates particle filtering with Markov Chain Monte Carlo (MCMC) sampling to estimate the posterior distribution over model parameters. Like previous approaches, HOPE selects stimuli by maximizing expected information gain. Unlike typical grid based approaches, however, it overcomes the exponential scaling problem, which enables efficient and fast sampling of even high-dimensional parameter spaces to use in a behavioral experiment. HOPE makes it feasible to estimate multidimensional psychometric functions in many more dimensions than previously possible.

## 2 Results

### High-dimensional Online Particle Estimation (HOPE)

Consider a typical psychophysical experiment, in which a participant is asked either to classify a single stimulus into one of two categories (e.g., “A” vs. “B” or “signal” vs “noise”), or is presented with several stimuli and decides which one belongs to a category in question (e.g. category “A”). The stimuli are defined by a vector of feature values (e.g., discs that vary in size, color and motion direction would have three stimulus feature dimensions), and a psychometric function (see e.g. Wichmann and Hill, 2001) is used to define the relationship between the stimulus features and a response probability (Figure 1). This function contains a set of parameters describing the precise form of this relationship.

**Figure 1:**
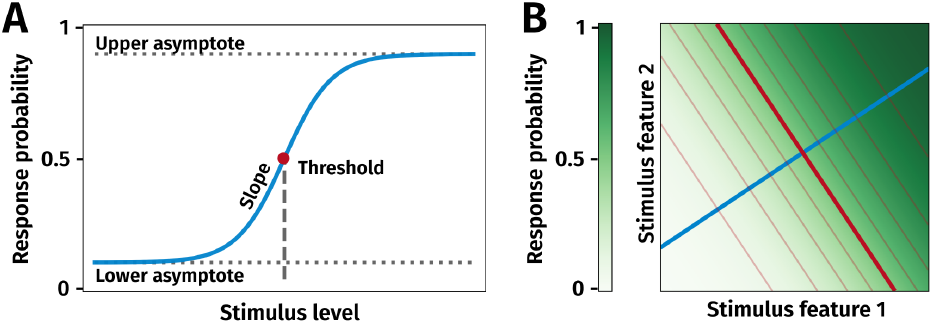
Psychometric functions in 1D (**A**) and 2D (**B**). The threshold in the one-dimensional psychometric function, visualized as the red dot in **A** becomes a line in two dimensions (red line in **B**). Its direction characterizes a hyperplane with no change in response probabilities. Any cut orthogonal to the red line (e.g. the blue line) can be imagined as the s-shaped curve from the one-dimensional psychometric function plot. The color gradient in **B** visualizes this s-shaped nonlinearity for the 2D space. The additional lines parallel to the red line are placed every 0.1 step of probability and visualize the s shape with larger changes in probability closer the threshold line.

The empirical goal is to estimate the most plausible psychometric function parameters from behavioral data. The experimenter’s current belief about the plausible parameter values is expressed as a probability distribution over the parameters, and each new stimulus-response pair (trial) is used to update this belief using Bayes’ Theorem. This posterior probability distribution is then used to guide stimulus selection for the next trial, by choosing the stimulus expected to most reduce the uncertainty of the posterior (or equivalently, to maximize expected information gain (Lindley, 1956; MacKay, 1992)). Posterior updates and the selection of the next stimulus must therefore be performed between consecutive trials, and fast enough that the computational cost does not outweigh the benefits of efficient stimulus selection. However, grid-based approximations do not scale to high dimensions, and commonly-used Markov Chain Monte Carlo (MCMC) algorithms (e.g. Geman and Geman, 1984; Duane et al., 1987; Hoffman and Gelman, 2014) are too slow for online adaptive data collection as incorporating new observations typically requires restarting the sampling procedure from scratch.

Pairing particle-filtering with MCMC steps (Kujala and Lukka, 2006) overcomes these limitations. The posterior is approximated using a finite set of samples, called particles, each representing a possible parameter vector of the high-dimensional psychometric function (Figure 2A). These particles are initially distributed according to the prior distribution. After each new response, particles are re-weighted according to the likelihood of the observed data under the parameter vector represented by the particle, then resampled proportionally to their weight (Figure 2B), keeping the number of particles the same. Over iterations, the particles concentrate in regions of higher posterior density; low probability particles are likely to disappear. However, this algorithm alone leads to particle degeneracy, in which an increasing fraction of the posterior mass becomes concentrated on a small number of particles, reducing the representational capacity of the particle set and impairing inference. To prevent this, the position of each particle is slightly perturbed via several MCMC steps (Figure 2B). The updated particle set approximates the posterior given all trials observed so far.

**Figure 2:**
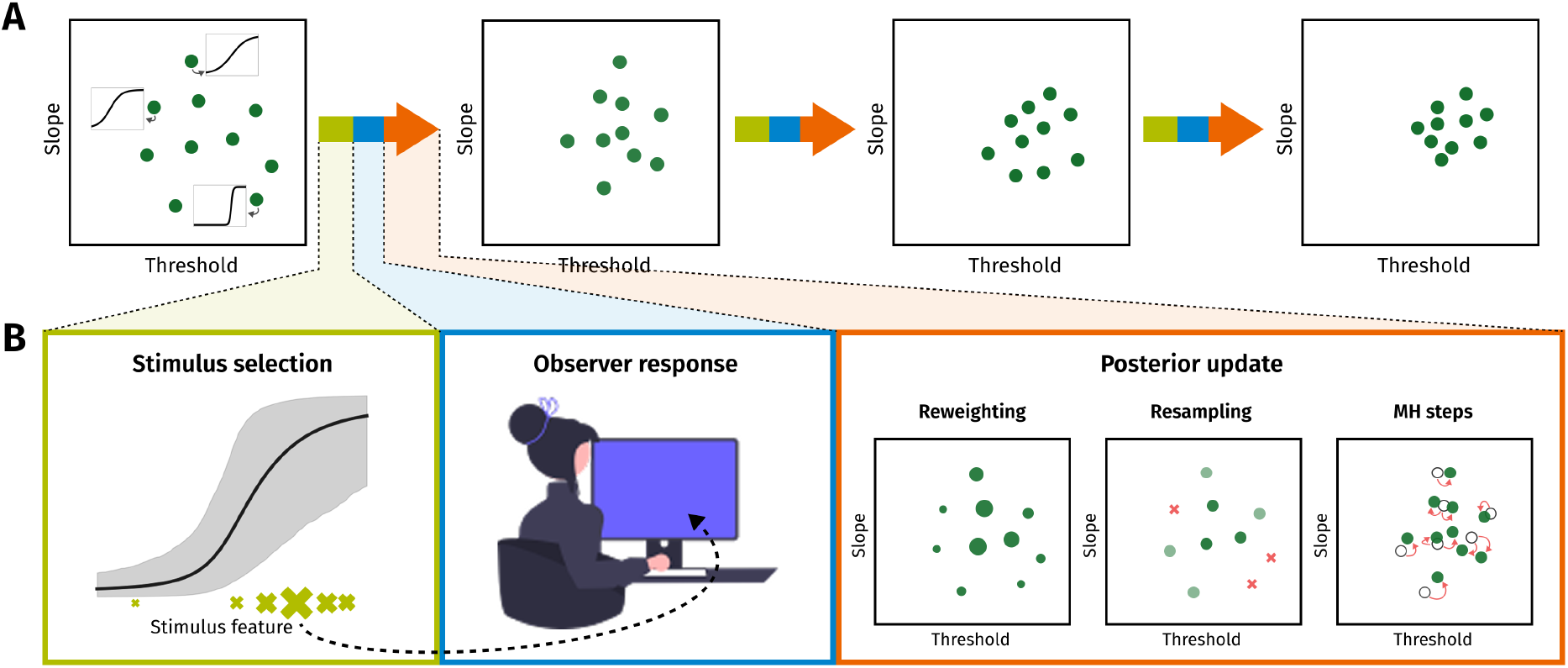
Running an adaptive experiment. **A**: Example progression of particles over several trials. The x- and y-axes represent two parameters of a one-dimensional psychometric function, the slope and threshold. The small figures in the first box show what the functions corresponding to the parameter combinations might look like. From left to right new data is collected and the posterior entropy decreases, as indicated by the particles moving closer together. **B**: Steps between one posterior and the next. First, a new stimulus is selected based on the expected information gain. Here, the gray area represents posterior uncertainty and the size of the stimuli at the bottom (green crosses) indicates the expected information gain from selecting the respective stimulus. Then, the selected stimulus is presented to the observer and a response is recorded. Finally, the posterior is updated based on the new data point. The size of circles in the reweighting box represents the weight for each particle. In the resampling the center three points are sampled twice (darker green), while three other points with low weights are not sampled at all (red crosses). In the MH steps, each particle is moved slightly to more effectively represent the posterior distribution.

Once the posterior has been updated, HOPE selects the next stimulus from a discrete set of stimuli. Uncertainty is quantified as the entropy of the posterior distribution: distributions with more diffuse probability mass have more uncertainty about plausible parameter values and thus higher entropy. The algorithm uses the current set of particles to simulate possible observer responses for each candidate stimulus, then chooses the stimulus that is expected to lead to the largest reduction in entropy. Critically, the same set of particles is carried forward and refined throughout the experiment, and both posterior updating and stimulus selection operate directly on this fixed set of particles, meaning that computation remains fast enough for real-time use in behavioral experiments.

### Simulations

We first evaluated HOPE in simulations to assess both accuracy and efficiency in recovering the parameters of a multidimensional psychometric function, relative to known ground truth values. To match the dimensionality of our behavioral experiment (see below), we define an 18-dimensional logistic psychometric function, with a 15-dimensional stimulus space (one weight per stimulus feature; the weight vector determines the orientation and slope of the plane around the decision boundary; Figure 1B), a bias term, and two lapse-rate parameters to capture stimulus-independent errors. Importantly, the number of parameters grows linearly with the number of stimulus features. Performance was compared to a baseline that sampled stimuli uniformly at random from a predefined pool, as no other method can currently perform adaptive sampling in 18 dimensions. Simulations were repeated 15 times, each time with a newly sampled virtual observer that was used in one baseline and one adaptive run, respectively.

Across simulations, the entropy of the posterior decreased more rapidly with HOPE than with the baseline (Figure 3A), while maintaining higher accuracy in recovering ground truth parameters (Figure 3B). For analyzing our data after simulation we used NumPyro (Bingham et al., 2019; Phan et al., 2019) with NUTS (Hoffman and Gelman, 2014) to compute the posterior for the psychometric function model from the relevant (adaptive or baseline) data, and used the posterior mean as the model for analysis. Accuracy was measured as the root mean squared error (RMSE) between the true parameters and those inferred by each method, and uncertainty in the parameter estimates was tracked using posterior entropy. For example, the adaptive method reached the same parameter recovery accuracy in roughly half the number of trials as the baseline (3,000 compared to 6,000). Although the adaptive method exhibited greater variability in RMSE (Figure 3B), even its maximum errors remained within the baseline’s standard deviation. Additionally, for each baseline-adaptive observer pair, the adaptive observer’s RMSE was always below the baseline observer. Therefore, HOPE not only matches but exceeds the baseline in accuracy, while substantially reducing the number of trials needed for reliable parameter estimation.

**Figure 3:**
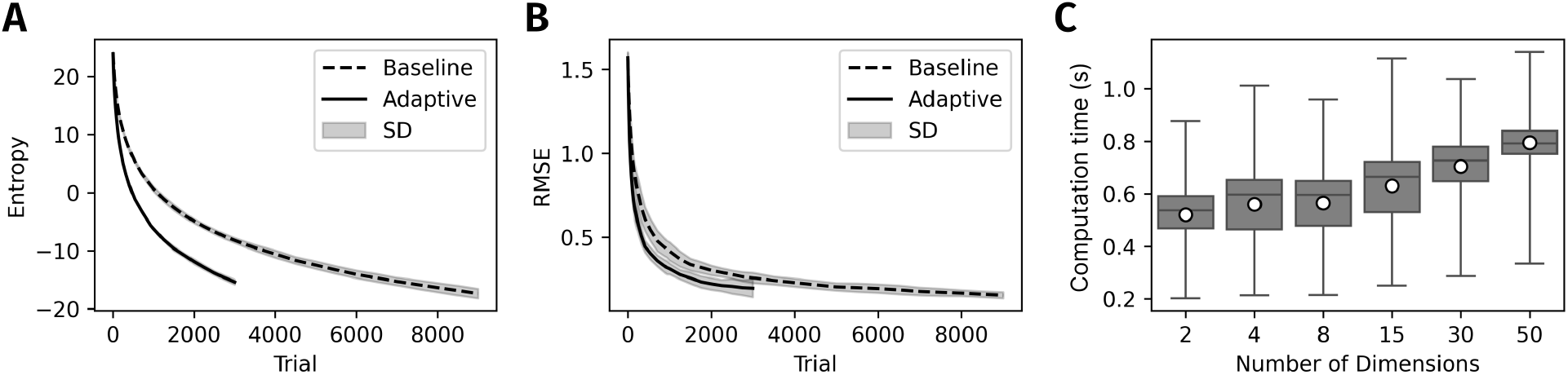
Results of simulated experiments. **A** and **B**: Experiments with psychometric function for 15-dimensional stimulus feature space. The lines represent the mean over 15 runs, respectively, the gray areas display the standard deviation. **A**: The posterior entropy (uncertainty) decreases faster for the adaptive experiment. **B**: The RMSE between ground truth and fitted parameters decreases faster if stimuli are sampled adaptively. **C**: Computation times for the posterior update plus stimulus selection for different posterior dimensionalities. The whiskers indicate the full range of observed times, while the edges of the box correspond to the 25th and 75th percentiles. The central line denotes the median, and the superimposed point represents the mean. Even for a 50-dimensional model parameter space, the time never exceeds 1.2 seconds.

We also evaluated the computation time of HOPE, finding that the entire posterior update and stimulus selection loop remained practical up to at least 50 parameter dimensions. Figure 3C shows the time required to select a new stimulus across dimensions ranging from 2 to 50, averaged over 15 simulations and 1000 trials, for a stimulus pool of 10,000 examples and 1,000 particles, on a standard computer. Even for 50 dimensional parameter spaces, the algorithm was able to provide the next stimulus in under a second for almost all trials. The parameters for these simulations were chosen to match our behavioral experiment (see below). While the stimulus set size and particle number will also influence computation time, we do think these are reasonable values that could be used for other experiments.

### Experiment: Perceived gender categorization in humans

We next evaluated HOPE in a behavioral experiment. Participants classified face images generated via a 15-dimensional parametric face space (see Methods) as male or female, with stimuli selected from a fixed pool of 10,000 artificial faces using either HOPE or the baseline. Five participants completed a total of 12,000 trials across eight sessions, divided into 9,000 trials using baseline sampling and 3,000 trials using HOPE. As above, final posterior distributions were estimated using a standard MCMC procedure for analysis.

Consistent with the simulation results, HOPE achieved higher certainty about psychometric parameters in fewer trials. Posterior entropy decreased more rapidly under HOPE than under the baseline (Figure 4A). In fact, for four out of five of the participants, the baseline method did not reach the entropy of HOPE even within the 9000 baseline trials.

**Figure 4:**
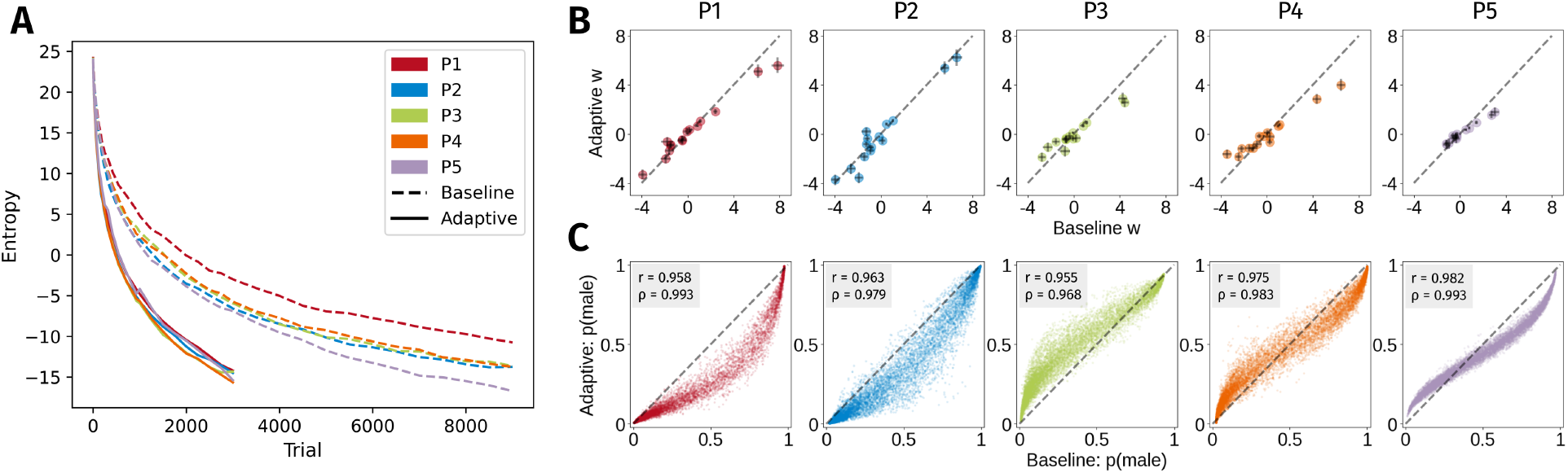
Results from a perceived gender categorization experiment with five participants. **A**: Entropy over trials for each participant. As in the simulations, the entropy decreases faster for adaptive than for baseline sampling. **B**: Model parameters for the 15 stimulus feature dimensions from the baseline (x-axis) versus adaptive (y-axis) fitted models for each participant. Error bars represent the 95% credible intervals. Values on the diagonal indicate agreement of the models.**C**: Predicted probabilities from baseline (x-axis) versus adaptive (y-axis) fitted models for each stimulus in the stimulus pool and for each participant. Close alignment with the diagonal indicates high agreement of the models.

While the entropy results were qualitatively similar in our real experiment as in the simulations, the next important question for the real experiment is whether the adaptive and baseline methods yielded similar decision functions. Because the true psychometric functions are unknown, we cannot directly quantify estimation accuracy as in the simulations. Instead, we measured the agreement between the adaptive and baseline methods. First, we directly compared the fitted weights of each participant’s MCMC posterior (Figure 4B). The close alignment with the diagonal indicates that both sampling strategies lead to similar estimates about the importance of the 15 features for each participant’s decisions. Second, we used the mean posterior model to compute the predicted probability *p* (male) for each stimulus in the dataset and compared them. Although the predictions were not identical, they showed a high degree of agreement (Figure 4C), with most points falling in the lower-left or upper-right of the plot, indicating strong agreement on stimuli perceived as clearly female or male. Predicted probabilities for intermediate cases clustered around the diagonal with high linear correlation values, forming a clear monotonic trend. Across participants, both the linear Pearson *r* and the rank order Spearman *ρ* correlation coefficients were consistently high with a minimum of 0.955 for r and 0.968 for *ρ* (see individual plots in 4C for each participant’s values).

To further evaluate the similarity of the results between the baseline and adaptive methods, we visualized the perceived continuum from female to male faces. Starting from a reference face (the mean of the original face dataset), we generated example stimuli based on the psychometric function of the average baseline and adaptive observer, by moving along the direction of the mean weight vector. Since this vector represents the direction with maximum female-male differentiation, we call this the decision vector (e.g. orthogonal to the blue line in Figure 1B), similar to the “decision faces” in Macke and Wichmann (2010). We selected points along this vector corresponding to predicted classifications from very female to very male. Viewing the resulting face stimuli (Figure 5A) yields the subjective impression that the discovered decision vectors indeed capture information about perceived gender when the results are averaged over participants. At the same time the resulting faces from the average baseline and adaptive observer appear similar, indicating that both methods recover similar decision functions.

**Figure 5:**
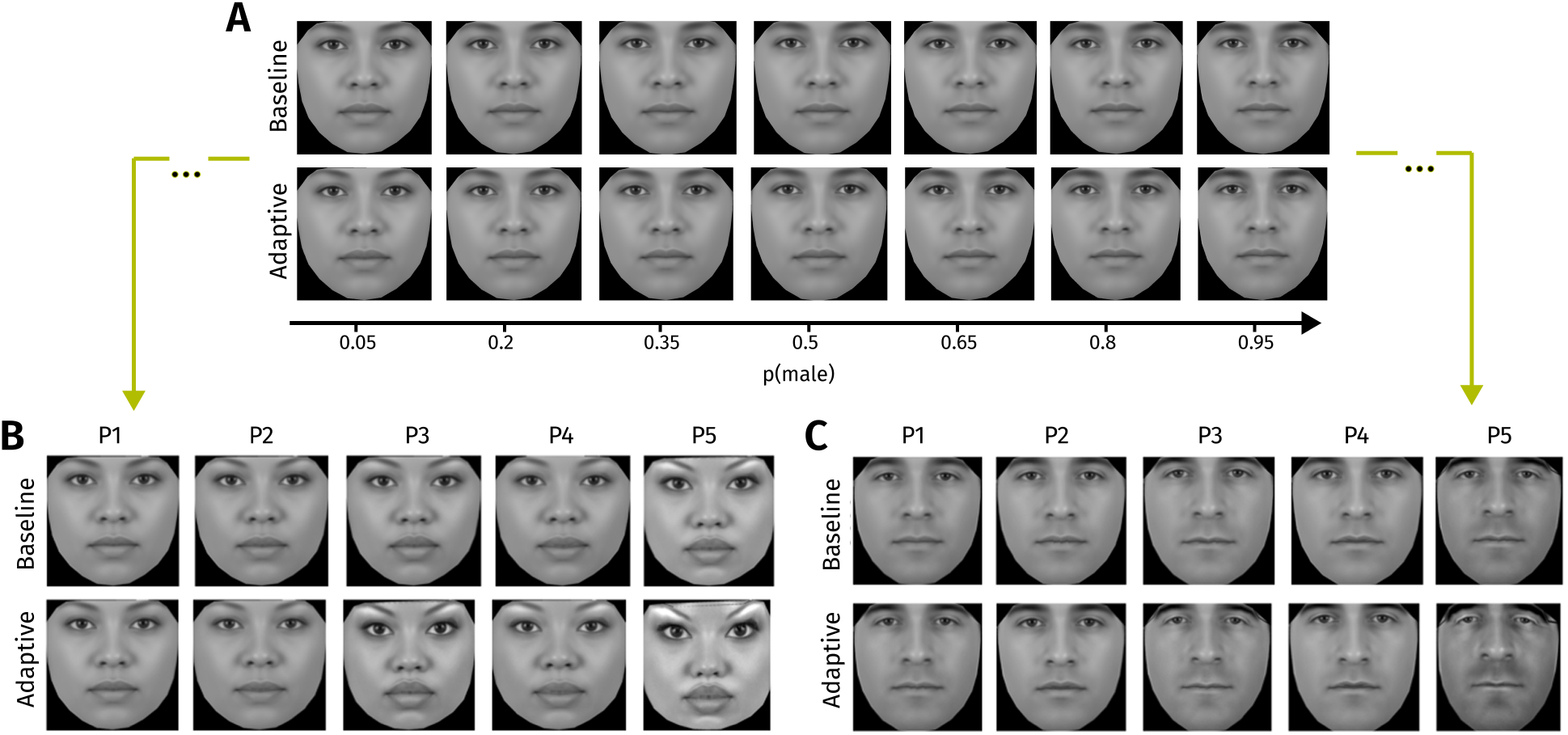
Visualization of participants’ female-male decision direction. Starting from a common reference face (mean of the face dataset (Ma et al., 2015)), we generated faces along the decision vector to show the continuum from female to male according to different estimated models. **A** shows the mean participant’s continuum for the baseline (9000 trials, top) and adaptive (3000 trials, bottom) sampling procedure, which appear highly similar, indicating the methods recover similar decision functions. In **B** and **C** we moved along each participant’s decision vector to generate highly female (**B**) and highly male (**C**) facial exemplars. Note that these are extreme faces for visualization purposes; similar faces were not shown in the experiment. As in the experiment, all of the faces here were generated from a point in our 15-dimensional face feature space and do not belong to any real person. Overall, faces generated from different model fits within the same participant (columns) appear more similar than to faces generated across participants (rows), indicating that participant-specific features were reliably recovered using both baseline and adaptive sampling.

Finally, we assessed whether the adaptive and baseline methods qualitatively captured similar decision behavior within individuals. Similar to the continuum in Figure 5A, we generated very female (Figure 5B) and male (Figure 5C) faces according to each participant’s psychometric function in the baseline and adaptive methods. This time we moved further along the decision vector, towards extreme female and male regions, to visually emphasize the differences between features. Subjectively, faces appear more similar between methods for the same participant than across participants, indicating that both approaches recover consistent and participant-specific representations. Together, the face visualizations suggest that the data collected with HOPE captures genuine perceptual structure rather than methodological artifacts, while preserving sufficient resolution to reveal individual differences in categorization.

## 3 Discussion

In this work we describe High-dimensional Online Particle Estimation (HOPE), which enables adaptive psychophysics experiments in high-dimensional feature spaces for parametric psychometric functions. We demonstrated its performance in simulations and in a perceived gender categorization experiment. Both in simulations and an experiment, our adaptive stimulus selection reduced parameter uncertainty faster than a random sample baseline. This improvement enables shorter and more efficient psychophysics experiments in practice. To spread HOPE, we make our software available as an open source software package (Reining and Turon, 2026).

We make two key contributions: first, we reimplement the method of Kujala and Lukka (2006), which uses a more efficient and robust posterior update than other methods. Incorporating new data into standard MCMC methods typically requires restarting sampling, meaning that inference is on the order of minutes or even hours rather than seconds for any sizable dataset. Particle filtering solves this issue. DiMattina (2015b) proposed a particle-filtering approach that is similar to HOPE, demonstrating it for four stimulus features. However, this method does not include the Metropolis-Hastings steps to avoid particle degeneracy (Kujala and Lukka, 2006), which can lead to poor approximation of the true posterior.

Second, we extend this basic algorithm to handle stimulus spaces of substantially higher dimensions than previously possible, fast enough to be used in behavioral experiments in a standard trial loop, and demonstrate empirically that this works. Methods that use grid approximation (e.g. Watson and Pelli, 1983; Watson, 2017; Kontsevich and Tyler, 1999; Vul et al., 2010) are limited to four or five parameter dimensions. The non-parametric AEPsych (Owen et al., 2021, see below) is currently only feasible for approximately eight stimulus features. Two related approaches that can be applied to high-dimensional stimuli use MCMC algorithms directly with people’s categorical or adjustment responses (Harrison et al., 2020; Sanborn and Griffiths, 2007). However, the goal of these methods is to estimate the subjective probability distributions that people associate with different categories, not to explicitly model the functional relationship between stimulus features and categorical responses.

One open issue with the use of any adaptive method is the possibility that experiencing adaptively-selected stimulus sequences may itself cause changes in the participants’ decision functions. We investigated short-term sequential dependencies in our human data, and found that responses in blocks using adaptive stimulus selection did indeed show larger contributions of a history term to the variance of the linear prediction term inside the logistic function. However, the history contribution to the variance stays small (for example, for a history of one response, the history contribution to variance was estimated to be less than 7% for all participants, compared to the stimulus-related variance of 93%). On top of this, we think that the higher relative contribution of the history term to the variance in the adaptive condition could be explained by a lower variance of stimulus-related variance due to the fact that the adaptive procedure selects more stimuli with a similar probability range (for more details see Supplementary Material). While for our human experiment, adaptive and non-adaptive experiments led to largely similar perceived gender decision functions (Figure 4 and 5), experimenters should be aware of this caveat when using adaptive stimulus selection.

A major practical advantage of HOPE is its flexibility: it can be combined with arbitrary parametric forms describing the purported decision function. Although we employ a logistic model with a linear predictor in the present work, other sigmoidal distributions, models incorporating nonlinear interactions among stimulus features, or indeed non-sigmoidal or non-monotonic parametric functions linking stimulus properties to response probabilities should all be theoretically possible to estimate. Currently however, HOPE is limited to discrete response alternatives. While computing the posterior distribution would be possible for multinomial or even continuous response distributions, efficiently computing the expected information gain over continuous responses to select the next stimulus is much harder.

Imposing parametric structure is particularly advantageous in high-dimensional stimulus spaces. In the absence of parametric assumptions, learning the psychometric function requires dense sampling across the stimulus space in order to capture local variations, which quickly becomes infeasible as dimensionality increases. In contrast, parametric models impose a global structure on the mapping from stimulus features to response probabilities, such that the number of free parameters typically grows much more slowly than the number of possible stimuli, linearly in the case of the linear predictor considered in our experiments. As a result, data collected at a limited set of stimulus locations can inform the psychometric function throughout the stimulus space, improving sampling efficiency. Indeed, a critical first step is the choice of a stimulus feature space that captures task-relevant aspects: different representations can make interactions between stimulus features more or less tractable, and finding a meaningful representation is often a non-trivial problem (Guan et al., 2018; Ryali et al., 2020b). Here, we have assumed that such a representation has already been specified (see Ryali et al. (2020a); Lind and Yu (2024) for more details on the representational space we used).

Though the assumptions inherent in parametric modelling can be useful or even necessary in high-dimensional stimulus spaces, they come with the concern that these assumptions may be violated in practice. How does HOPE perform when the parametric model is not an accurate description of behavior? When deviations between the true underlying psychometric function and the assumed parametric form are modest (e.g. data are generated from a different link function than the assumed logistic link), adaptive sampling remains effective, as it continues to concentrate measurements in informative regions of the stimulus space (see Supplementary Material). However, when the assumed parametric form is severely misspecified or even unknown, the adaptive procedure may bias stimulus selection toward regions that are uninformative for the true data-generating process, potentially rendering it less efficient than non-adaptive procedures that sample more broadly. Nonparametric approaches mitigate this by using even weaker assumptions about how stimuli and responses are related. A recent example is AEPsych (Owen et al., 2021), which uses Gaussian Process models to represent the psychometric surface. Currently, the cost of Gaussian process inference and acquisition optimization can limit real-time use unless approximate or sparse models are employed (see an application of this in Hong et al. (2026)). Nevertheless, the judicious pairing of parametric and non-parametric adaptive algorithms will widen the scope of behavioral experiments that can now be performed.

A number of experimental paradigms could benefit from HOPE, particularly those involving high-dimensional stimulus representations and limited trial budgets, for which reasonable parametric assumptions can be made. One example is cue combination experiments, in which multiple stimulus features jointly influence perceptual judgments and the goal is to characterize how these cues are weighted and integrated (e.g. Ernst and Banks, 2002; Alais and Burr, 2019; Landy et al., 1991). In such settings, the number of possible stimulus configurations grows rapidly with the number of cues, making exhaustive sampling inefficient. By concentrating measurements in regions that are most informative about cue weights and interactions, the present method could substantially reduce the number of trials required to estimate these quantities. Similarly, another potential application is in empirically estimating high-dimensional contrast sensitivity functions, which can characterize visibility across stimulus features such as spatial and temporal frequency, size, eccentricity, luminance and chromaticity (Mantiuk et al., 2022; Ashraf et al., 2024). Finally, the method could also be applied to reverse correlation paradigms, where stimuli are constructed from many elementary features (e.g., pixels or basis functions) and the objective is to infer the features that drive perceptual decisions or neural responses (e.g. Neri and Heeger, 2002; Murray, 2011; Park and Pillow, 2012). When the dimensionality of the feature space is moderate, parametric models combined with adaptive sampling may provide a more efficient alternative to traditional random sampling approaches, enabling faster recovery of decision-relevant structure. More broadly, HOPE may be useful in any decision making setting that matches the constraints outlined above and where exhaustive sampling of the stimulus space is infeasible.

## 4 Methods

Additional description of the Materials and Methods can be found in the Supplementary Material.

### High-dimensional Online Particle Estimation

In all simulations (across different stimulus feature dimensionalities) and in the human experiment, we map the *k*-dimensional stimulus feature space onto a binary response using a multidimensional psychometric function

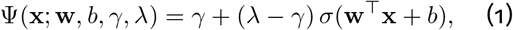

where **x** is the *k*-dimensional stimulus feature vector, *b* is a bias term, **w** is a *k* dimensional weight vector, *γ* and *λ* are the lower and upper asymptotes, respectively (reflecting stimulus-independent lapses) and 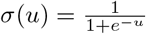 is the logistic sigmoid function.

Denoting the parameters of the psychometric function collectively as *θ*, the first goal of HOPE is to estimate the posterior distribution given the data observed so far. The data {(**x**_1_, *y*_1_), …, (**x**_*n*_, *y*_*n*_)} consist of stimuli **x**_1:*n*_ = {**x**_1_, …, **x**_*n*_} and the corresponding responses *y*_1:*n*_ = {*y*_1_, …, *y*_*n*_}, after *n* trials. Denoting the likelihood as *p*(*y*_*n*_ | **x**_*n*_, *θ*), with *p*(*y*_*n*_ = 1 | **x**_*n*_, *θ*) = Ψ(**x**_*n*_; *θ*) and *p*(*y*_*n*_ = 0 | **x**_*n*_, *θ*) = 1 − Ψ(**x**_*n*_; *θ*), we seek to compute the posterior distribution over the parameters

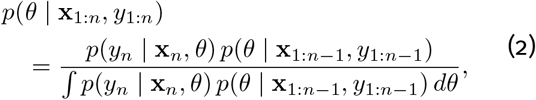

which is analytically intractable. The particle filtering approximation at trial *n* consists of a set of *m* samples {*θ*_1,*n*_, *θ*_2,*n*_, …, *θ*_*m,n*_}, called particles. When new data is added, the particles are updated in three steps:

1. *Re-weighting*: The current posterior *p*(*θ* **x**_1:*n*_, *y*_1:*n*_), represented by the *m* particles {*θ*_1_, |*θ*_2_, …, *θ*_*m*_}, assigns equal weight to all of its particles. We assign new weights *β*_*i*_ based on the likelihood of our new data point **x**_*n*+1_, *y*_*n*+1_ and normalize with the sum over all weights:

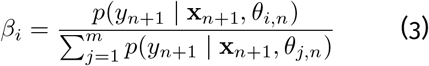
2. *Resampling*: Using the assigned weights we now sample *m* times from the particles {*θ*_1_, …, *θ*_*m*_} with replacement, i.e. *p* (*θ*_*i*_) = *β*_*i*_. Therefore, particles with higher likelihoods given the data are more likely to be sampled.
3. *Metropolis-Hastings*: Since the positions of the particles are fixed in the previous two steps, resampling typically leaves us with a large proportion of duplicate particles, particularly when new data rule out much of the region the particles previously occupied. To re-establish a representative approximation to the posterior, we let each particle take a few steps to diversify their locations. For this we use the Metropolis-Hastings algorithm (Metropolis et al., 1953), which is commonly used in the proposal and update step of MCMC. For each particle we perform *n*_MH_ Metropolis-Hastings steps (see Supplementary Methods for full description). As an additional benefit, MH steps move particles toward regions of higher probability, which helps keep the particle approximation concentrated in regions of high probability mass over time.

Using this approximation to the posterior, we can now select the stimulus **x**_*n*_ for the current trial by choosing the stimulus to minimize uncertainty about the model parameters (Kontsevich and Tyler, 1999). For a multidimensional posterior, uncertainty is quantified by the entropy *H*(Θ), where Θ denotes the random variable over parameter values *θ*. Selecting the next stimulus to minimize posterior entropy is equivalent to maximizing the reduction in entropy (or, alternatively, the information we gain) induced by the next observation under that stimulus (Kujala and Lukka, 2006; Lindley, 1956; MacKay, 1992):

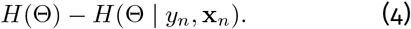

Because the response *y*_*n*_ to a candidate stimulus **x**_*n*_ is not yet known, we maximize the *expected* entropy reduction over possible responses, which is equivalent to subtracting the conditional entropy *H*(Θ | *Y*_*n*_, **x**_*n*_) over both Θ and *Y*_*n*_ in equation 20. This yields the mutual information between parameters and response, conditioned on the candidate stimulus,

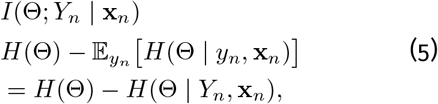

which quantifies how much, on average, observing *Y*_*n*_ will reduce uncertainty about Θ. Using the symmetry of mutual information (Lindley, 1956), this objective can equivalently be written as

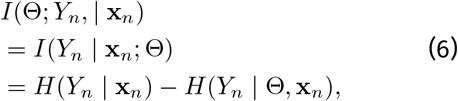

where both terms depend on the candidate stimulus **x**_*n*_. Here, *H*(*Y*_*n*_ | **x**_*n*_) is the entropy of the predictive distribution *p*(*y*_*n*_ | **x**_1:*n*_, *y*_1:*n−*1_), reflecting the total predictive uncertainty about the upcoming response, whereas *H*(*Y*_*n*_ | Θ, **x**_*n*_) is the expected response entropy given the parameters, reflecting the component of that uncertainty that is not attributable to uncertainty about the parameters. This reformulation already used by Kujala and Lukka (2006) is computationally convenient under our particle approximation of the posterior. The entropy of the predictive distribution can be computed from our particles (Kujala and Lukka, 2006) as:

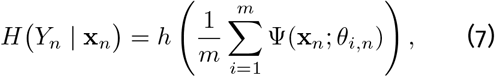

where *h*(*p*) = −*p* log −*p* (1 − *p*) log(1 − *p*) is the binary entropy function. Thus, this term corresponds to the binary entropy of the average predicted response probability. The conditional entropy is given by the average binary entropy under each particle,

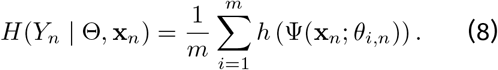

Finally, we compute the expected information gain for any candidate stimulus **x**, and select the stimulus that maximizes it:

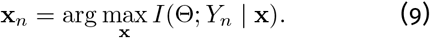

In practice, this objective is evaluated over a set of candidate stimuli, either from a predefined pool or drawn from a stimulus space.

### Stimulus pool

We applied the Active Appearance Model (AAM) to the neutral, grayscale faces of the Chicago Face Database (CFD) (Ma et al., 2015; Guan et al., 2018). The resulting model contains both shape and texture features, providing a good starting point for features that could be important for perceived gender categorization. We then performed a PCA and used the first 15 PCs to obtain a face feature space that contains 87.1 % of the variability and can easily be used for optimization during our experiment.

The CFD contains 597 neutral faces of which 307 are female and 290 are male. To cover more of the space of faces in the 15 dimensions without crossing into unnatural areas of the space, we generated new stimuli as a linear combination of two faces from the CFD. Since we aimed to measure participant’s individual decision functions for differentiating between female and male, we created linear combinations of female and male faces only. For each sampled female-male face combination, we generated several morphs with different weight contributions of the female and male face. Our final stimulus pool created in this way contained 10,000 stimuli (see Supplementary Material for a more detailed description and examples).

We used this stimulus pool both in the simulations for the 15-dimensional feature space and in the experiment with human participants. We found that the adaptive algorithm often selected the same stimulus successively. Since faces are easily recognizable stimuli, this can have a strong effect on the independence of trials. Therefore, we decided to sample from the stimulus pool without replacement within each block of 100 trials.

### Simulations

We simulated artificial observers using the 15-dimensional psychometric function (Equation 1) by fitting the weights to the ground-truth CFD labels of a random 80 % subset of the CFD faces, and sampling *γ* and *λ* from a *Beta*(1, 30) and *Beta*(30, 1) distribution, respectively. We repeated this procedure 15 times to yield 15 independent and slightly different artificial observers. We then ran simulations where each artificial observer performed the perceived gender categorization task on the stimulus pool described above. In each trial *t*, the artificial observer received a stimulus feature vector **x**_*n*_, selected either adaptively (3,000 trials) or randomly (9,000 trials) and responded with a binary label *y*_*n*_ sampled from the Bernoulli distribution defined by the psychometric function with the groundtruth parameters of the artificial observer.

To estimate the stimulus selection time, we created artificial observers for each dimensionality *n* by sampling the coefficients of a n-dimensional logistic regression from a standard normal distribution and lower and upper lapse rates from *Beta*(1, 30) and *Beta*(30, 1) distributions. We then created an artificial stimulus pool of 10,000 stimuli by sampling uniformly from the n-dimensional hypercube [− 5, 5]^*n*^. For each dimensionality and artificial observer, we ran 1000 trials with HOPE and measured the time it took our algorithm to select each stimulus.

### Perceived gender categorization experiment

Five participants naive to the adaptive-baseline comparison were presented with face stimuli (subtending approximately 12 degrees of visual angle) for up to 1 s, and responded whether they perceived the gender of the face to be male or female via button press. No feedback was provided. Participants performed eight sessions of 1,500 trials each, using either adaptive (two sessions) or baseline (six sessions) sampling for the entire session; the order of sessions was randomized for each participant. In adaptive sessions, particles were initialized by sampling from the following priors: lower and upper lapse rates were assigned *Beta*(1, 30) and *Beta*(30, 1) priors, respectively, while the psychometric function coefficients and bias term were assigned standard normal (*N* (0, 1)) priors. All study procedures were approved by the Ethics Committee of TU Darmstadt (EK-77/2022). Further details can be found in the Supplementary Methods.

## 5 Data, Materials and Software Availability

The code and data used to produce the results reported in this study are available at https://doi.org/10.5281/zenodo.21098788.

## 6 Acknowledgements

This work was supported in part by the Hessian Ministry of Higher Education, Research, Science and the Arts and its LOEWE research priority program ‘WhiteBox’ under grant LOEWE/2/13/519/03/06.001(0010)/77, by the Deutsche Forschungsgemeinschaft (German Research Foundation, DFG) under Germany’s Excellence Strategy (EXC 3066/1 “The Adaptive Mind”, Project No. 533717223), and by the European Union (ERC, SEGMENT, 101086774). Views and opinions expressed are however those of the author(s) only and do not necessarily reflect those of the European Union or the European Research Council. Neither the European Union nor the granting authority can be held responsible for them.

## 7 Author contributions

**Conceptualization**: RT, LCR, PAH, FJ, and TSAW.

**Data curation**: LCR and PAH.

**Formal analysis**: RT, LCR, and PAH.

**Funding acquisition**: FJ and TSAW.

**Investigation**: RT, LCR, PAH, and TSAW.

**Methodology**: RT, LCR, PAH, CL, AJY, FJ, and TSAW.

**Project administration**: RT and LCR.

**Resources**: FJ and TSAW.

**Software**: LCR, PAH, CL, and AJY.

**Supervision**: FJ and TSAW.

**Validation**: RT, LCR, and PAH.

**Visualization**: RT, LCR, and TSAW.

**Writing - original draft**: RT.

**Writing - review & editing**: RT, LCR, LS, AJY, CAR, FJ, and TSAW.

## Appendix

Here we provide a more detailed and complete description of all methods from the main paper.

### High-dimensional Online Particle Estimation

#### Parametric model

As given in the main paper, we use a multidimensional psychometric function, parameterized as:

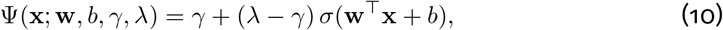

where **w** and **x** are k-dimensional. With this parameterization the decision boundary is a (*k* −1)-dimensional hyperplane parameterized by a weight vector **w**, which we also call the decision vector and which is orthogonal to the plane, and a bias term *b*, which shifts the boundary along this orthogonal direction. The classical psychometric function parameters, slope and threshold, are jointly encoded in these parameters: the norm ∥**w**∥ determines how rapidly response probabilities change with distance from the decision boundary, while the bias *b* determines the location of the boundary corresponding to the midpoint of the psychometric function. More generally, planes parallel to the decision boundary correspond to iso-probability surfaces.

To account for lapses, the psychometric function includes a lower asymptote *γ* and an upper asymptote *λ*, reflecting that observers may respond incorrectly even for very easy stimuli. In the simulations shown in Fig. 3A,B and in the human experiment, the stimulus dimensionality is fixed at *k* = 15; in Fig. 3C, *k* varies as indicated on the x-axis.

We refer to the parameters of the psychometric function, {**w**, *b, γ, λ*, λ}, collectively, as *θ*. The psychometric function Ψ(**x**; *θ*) gives the probability for a response *y* of 1. In turn, the probability to respond with *y* = 0 is given by 1 − Ψ(**x**; *θ*).

#### Bayesian framework

{} {}

{}

In our experiment we aim to estimate the underlying parameters of the model describing the generative decision process of each observer. For this, we assume the multidimensional psychometric function Ψ from above, and infer the underlying parameter values from the data we collect. The main text introduces the posterior update rule for the psychometric function parameters. Here we derive this update in full and discuss how it can be approximated in practice. Given the observer’s data {(**x**_1_, *y*_1_), …, (**x**_*n*_, *y*_*n*_)}, consisting of stimuli **x**_1:*n*_ = {**x**_1_, …, **x**_*n*_} and the corresponding responses *y*_1:*n*_ = {*y*_1_, …, *y*_*n*_}, after *n* trials, Bayes’ theorem tells us what parameter combinations are more or less probable with the posterior distribution

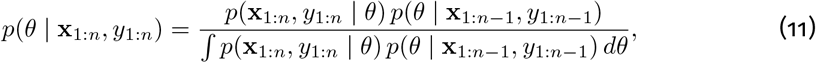

where *p*(*θ* | **x**_1:*n−*1_, *y*_1:*n−*1_) is the posterior after the previousJ trial, acting as a prior in this trial, *p*(**x**_1:*n*_, *y*_1:*n*_ | *θ*) is the likelihood function and *p*(**x**_1:*n*_, *y*_1:*n*_ | *θ*^*′*^) *p*(*θ*^*′*^ | **x**_1:*n−*1_, *y*_1:*n−*1_) *dθ*^*′*^ is a normalization constant. Since the trials are assumed to be independent and the previous posterior contains all information given the previous data, the likelihood can be rewritten as *p*(*y*_*n*_ | **x**_*n*_, *θ*) and we end up with

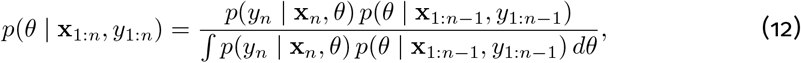

To compute the likelihood of an observed response *y*_*n*_ given a stimulus **x**_*n*_ and *θ*, we use the psychometric function from above with

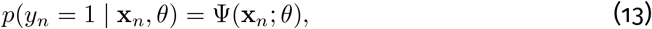

and

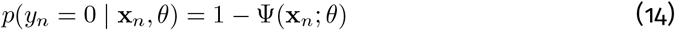

While equation 12 provides a theoretical solution to compute a posterior from a previous posterior or a prior and the likelihood, this can only be computed analytically in a few cases, for example if the prior or previous posterior and likelihood are conjugate. In our case, where the likelihood is a Bernoulli variable over a logistic function (the psychometric function), there is no closed form or analytical solution. Therefore, the posterior has to be approximated numerically. The following section explains what this numerical approximation could look like and how the approximated posterior is updated over time.

#### Posterior update

For approximations of continuous distributions that are not constrained to a specific form, like e.g. a Gaussian, there exist two common approaches: first, in grid approximation, the continuous distribution is evaluated at a predefined grid of locations. When normalized to sum to 1, these values then form a natural discrete approximation of a continuous distribution. The second approach also represents the continuous distribution with a discrete approximation, but instead of *evaluating* the distribution at fixed locations, the distribution is represented by a set of samples that mimic the density. This means that there are more samples at regions with high density and less samples in regions with low density. The most common method here is Markov Chain Monte Carlo (MCMC). It works by starting a chain of samples at one sampled location and then proposing new sampling locations and accepting or rejecting them in a way that ensures the chain spends more time in regions where the probability is high and less time in regions with low probability. The accumulated samples then approximate the distribution. Both of these approaches can be used for computing posteriors, if the likelihood function can be computed. In both cases we use the fact that the denominator of Bayes’ theorem is just a normalization factor and we can simplify equation 12 if we are only interested in proportionality:

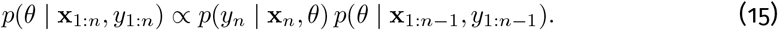

For grid approximation the normalization of a distribution is done by making sure that the sum of all grid values is one. This means that, when we have the values 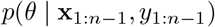 from the previous posterior for all grid locations we can simply multiply them with the likelihood given our new trial *p*(*y*_*n*_ | **x**_*n*_, *θ*) and make sure to normalize over the whole grid again. If the dimensionality of *θ* is low, this works well and is fast enough to update posteriors during an experiment (see e.g. Watson and Pelli, 1983). However, the grid grows exponentially with each dimension of *θ*, so with each dimension each posterior update needs exponentially more computations. Grid approximations are therefore too slow when *θ* contains more than four or five dimensions.

For approximating posteriors with MCMC, we start a chain with one sample *θ*_0_. This sample either comes from a prior or from the previous posterior. Then we run the chain: A proposal distribution proposes a new sample *θ*^*\**^ and we use some criterion based on the likelihood of our collected data and prior to decide whether we add this sample to our chain as *θ*_*i*+1_ or reject it and append *θ*_*i*_ to the chain. The proposal and acceptance step is crucial to ensure that the accumulated samples approximate the posterior distribution. The accumulated chain {*θ*_0_, *θ*_1_, …, *θ*_*m*_} then reflects the posterior distribution. In contrast to grid approximation, MCMC does not suffer from the same exponential growth in computation time with each added dimension to *θ*. However, MCMC cannot use much information from the previous posterior, but always starts a chain with just one sample and then has to be run long enough to approximate the posterior well. This process is normally computationally too slow to be used in real-time during an experiment.

Our solution, a particle filtering algorithm with Metropolis-Hastings updates (Metropolis et al., 1953), makes use of the previous posterior and does not suffer from the exponential growth with each dimension. Similar to MCMC, the posterior approximation at trial *n* consists of a set of samples {*θ*_0,*n*_, *θ*_1,*n*_, …, *θ*_*m,n*_}, which we call particles. However, when we collect new data, instead of starting the posterior computation from scratch, we update the posterior, i.e. our particles, from the previous step. This contains three steps:

1. *Re-weighting:* The current posterior *p*(*θ* | **x**_1:*n*_, *y*_1:*n*_), represented by the *m* particles *θ*_0_, *θ*_1_, …, *θ*_*m*_, assigns equal weight to all of its particles. We assign new weights *β*_*i*_ based on the likelihood of our new data point **x**_*n*+1_, *y*_*n*+1_ and normalize with the sum over all weights:

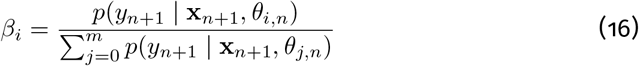
2. *Resampling*: Using the assigned weights we now sample *m* times from the particles {*θ*_1_, …, *θ*_*m*_ *}*with replacement, i.e. *p* (*θ*_*i*_) = *β*_*i*_. Therefore, particles with higher likelihoods given the data are more likely to be sampled.
3. *Metropolis-Hastings*: To diversify the particles, which now might contain duplicates, we use the Metropolis-Hastings algorithm, which is commonly used in the proposal and update step of MCMC. which is commonly used in the proposal and update step of MCMC. For each particle we perform *n*_MH_ Metropolis-Hastings steps, which work as follows:

Proposal: Draw a new proposal particle 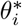 from a proposal distribution *q* around its original location:

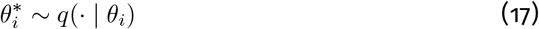

Acceptance probability: Compute the Metropolis-Hastings acceptance probability:

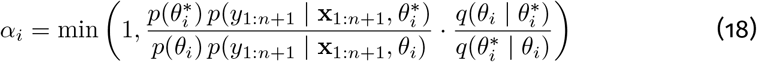

where *p*(*θ*) is the value of the prior evaluated at *θ*. Accept/Reject: Accept the proposed particle with probability *α*_*i*_:

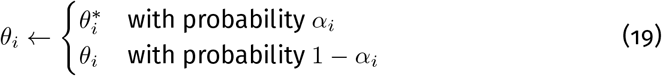

Re-weighting and resampling (also called importance resampling) alone leads to degeneracy: over time, the particle set collapses to fewer and fewer distinct locations, with multiple particles stacked at the same point. The subsequent Metropolis-Hastings steps counteract this by dispersing degenerate particles, ensuring that the density is well-represented locally around each resampled location. Intuitively, importance resampling maintains a good global approximation, since it starts from particles spread across different regions of the density and reweights them according to the new trial data, while the MH steps restore a good local approximation by spreading particles around each resampled location. In the limit of infinitely many MH steps, each particle would fully converge to the posterior, recovering a completely independent sample.

#### Minimizing expected entropy

As detailed in the main text, stimulus selection proceeds by maximizing the expected information gain about the model parameters (Kontsevich and Tyler, 1999). We reproduce the full derivation here for completeness. For a multidimensional posterior, uncertainty is quantified by the entropy *H*(Θ), where Θ denotes the random variable over parameter values *θ*. Selecting the next stimulus to minimize posterior entropy is equivalent to maximizing the reduction in entropy (or, alternatively, the information we gain) induced by the next observation under that stimulus (Kujala and Lukka, 2006; Lindley, 1956; MacKay, 1992):

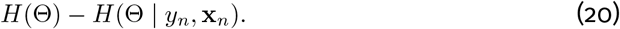

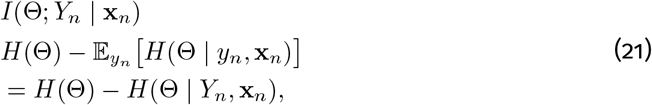

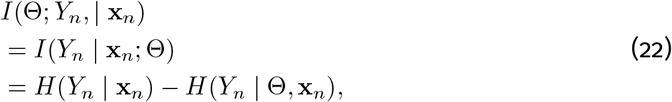

where both terms depend on the candidate stimulus **x**_*n*_. Here, *H*(*Y*_*n*_ | **x**_*n*_) is the entropy of the predictive distribution *p*(*y*_*n*_ | **x**_1:*n*_, *y*_1:*n−*1_), reflecting the total predictive uncertainty about the upcoming response, whereas *H*(*Y*_*n*_ | Θ, **x**_*n*_) is the expected response entropy given the parameters, reflecting the component of that uncertainty that is not attributable to uncertainty about the parameters. This reformulation, already used by Kujala and Lukka (2006) is computationally convenient under our particle approximation of the posterior. In particular, the entropy of the predictive distribution can be computed from our particles as:

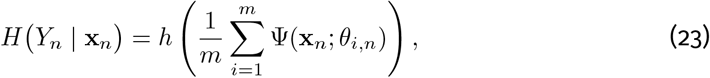

where *h*(*p*) = −*p* log *p*− (1 −*p*) log(1 − *p*) is the binary entropy function. Thus, this term corresponds to the binary entropy of the average predicted response probability. In contrast, the conditional entropy *H*(*Y*_*n*_ Θ, **x**_*n*_) is given by the average entropy under each particle,

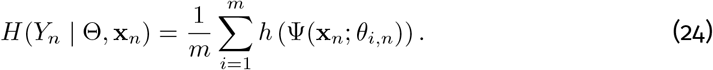

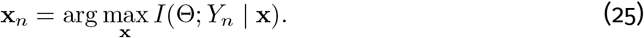

In practice, this objective is evaluated over a set of candidate stimuli, either from a predefined pool or drawn from a stimulus space. In our simulations and experiments, we used a predefined stimulus pool (see below).

### Stimulus pool for perceived gender categorization

Faces are complex stimuli in which just a few dimensions do not cover the whole range of variability and therefore are not enough to find differentiating features between different categories. Additionally, if one decides on too few features, this could bias participants into using these to differentiate between categories, even though other features could potentially be more important for the categorization that is investigated. At the same time, we propose an adaptive algorithm that is still limited in the number of dimensions it can be used on. To find a reasonable set of features, we performed the following steps. We applied the Active Appearance Model (AAM) to the neutral, grayscale of the Chicago Face Database (CFD) (Ma et al., 2015; Guan et al., 2018). The resulting model contains both shape and texture features, providing a good starting point for features that could be important for perceived gender categorization. We then performed a PCA and used the first 15 PCs to obtain a face feature space that contains 87.1 % of the variability and can easily be used for optimization during our experiment.

The CFD contains 597 neutral faces of which 307 are female and 290 are male. To cover more of the space of faces in the 15 dimensions without crossing into unnatural areas of the space, we generated new stimuli as a linear combination of two faces from the CFD. Since we aimed to measure participant’s individual decision functions for differentiating between female and male, we created linear combinations of female and male faces only. We sampled a female and male face from the CFD at random and then created their linear combination in the 15-dimensional space as *f*_*f*_ + *λ*\* (*f*_*m*_ − *f*_*f*_), where *f*_*f*_ is the 15-dimensional feature vector of the female face and *f*_*m*_ is the 15-dimensional feature vector of the male face. For each sampled pair we created the linear combinations for *λ* = {0.1, 0.2, …, 0.9} (note that the real faces at 0.0 and 1.0 were not part of the stimulus set). Our final stimulus pool created in this way contained 10,000 stimuli (see Figure 8 for examples).

### Simulations

The main text summarizes the simulation procedure used to evaluate parameter recovery. We give the full parameterization here. The goal of the simulations was to evaluate the performance of our adaptive algorithm in recovering known ground-truth parameters of a multidimensional psychometric function. For this we created artificial observers by fitting a 15-dimensional logistic regression to the ground truth labels of the CFD faces in the 15-dimensional stimulus feature space described above. To induce some variability, we repeated this procedure 15 times: for each repetition we fit a separate logistic model on a different random 80 % subset of the CFD faces. Additionally, for each artificial observer, we sampled lower and upper lapse rates from a *Beta*(1, 30) and *Beta*(30, 1) distribution, respectively. This results in 15 slightly different simulated observers in total.

We then ran simulations where each artificial observer performed the perceived gender categorization task on the stimulus pool described above. In each trial *t*, the artificial observer received a stimulus feature vector **x**_*n*_ and responded with a binary label *y*_*n*_ sampled from the Bernoulli distribution defined by the psychometric function with the ground-truth parameters of the artificial observer. We ran two versions of the simulation: one where stimuli were selected by our adaptive algorithm, and one where stimuli were selected uniformly at random from the stimulus pool (baseline). The simulation in the adaptive condition used the same priors as in the human experiment (see below). In the baseline condition, the simulations ran for 9,000 trials, while in the adaptive condition they ran for 3,000 trials.

#### Measuring stimulus selection time

The main text reports timing results across stimulus dimensionalities. The following describes how the corresponding artificial observers and stimulus pools were constructed. For each dimensionality *n* we created artificial observers by sampling the coefficients of a n-dimensional logistic regression from a standard normal distribution and lower and upper lapse rates from *Beta*(1, 30) and *Beta*(30, 1) distributions. We then created an artificial stimulus pool of 10,000 stimuli by sampling uniformly from the n-dimensional hypercube [−5, 5]^*n*^.

For each dimensionality and artificial observer, we ran 1000 trials with our adaptive algorithm and measured the time it took our algorithm to select each stimulus. We used the same priors as in the human experiment (see below).

### Perceived gender categorization experiment in humans

#### Participants

Five participants (2 female, aged 20 - 24 years) were recruited for the study. Participants were naive to the adaptive-baseline comparison of the study and just instructed to label presented faces as female or male. Participants received course credit for their participation. All study procedures were approved by the Ethics Commission of the TU Darmstadt (EK-77/2022).

#### Procedure

Participants performed the experiment in a dark room with blinds closed and additional black curtains. During trials participants rested their heads on a chin-rest with 55 cm distance to the screen.

The face categorization experiment consisted of eight sessions with 1,500 trials each. Each session used either adaptive or baseline sampling for the complete session, with two sessions using adaptive sampling and six baseline sampling. The order of adaptive and baseline sessions was assigned at random for each participant. In a baseline session each stimulus was sampled uniformly (with replacement) from the 10,000 stimuli in the stimulus pool. In an adaptive session the stimuli were selected from the stimulus pool with our adaptive algorithm. The particles were initialized by sampling from the following priors: lower and upper lapse rates were assigned *Beta*(1, 30) and *Beta*(30, 1) priors, respectively, while the psychometric function coefficients and bias term were assigned standard normal (*N* (0, 1)) priors. Each session was divided into blocks with 100 trials each, after which participants could take a break and continue when they were ready for the next block. In each block the adaptive algorithm sampled from the stimulus pool without replacement. After each block all stimuli were added to the stimulus pool again.

In each trial, participants performed a yes-no task with one face image presented on a black background for maximally one second. They reported whether they perceived this face as female or male with a button press on a button box. If participants responded during stimulus presentation, the presentation was cut short and they directly moved to the inter-stimulus-phase of one second, where a fixation cross was shown on a black background. If they did not respond during stimulus presentation, the next screen asked them whether they perceived the face as more male or more female. Once they responded, they moved to the inter-stimulus-phase and saw the black background with a fixation cross in the middle for one second.

Face stimuli could have slightly different sizes, because the AAM model allows different shapes for the faces, i.e. bigger or smaller in size overall or different width or height. Since the ratio and other features of the face shape are part of what could be used to perceive it as female or male, we did not force faces to fill a specific area on the screen. Additionally, the created face stimuli are generated on a black background, which is why the stimuli were shown on a black background as well to mask the transition from stimulus border to background. The stimuli were always placed centrally and extended between 12 and 13 degrees of visual angle.

#### Apparatus

All simulations, experiments, and analyses were conducted on a system running Ubuntu 20.04, equipped with a 13th Gen Intel Core i7-13700KF CPU and 32 GB of RAM. The computational processes used only CPU resources, with no GPU acceleration. We used Python 3.12.8 with key dependencies including PyTorch 2.5.1, NumPyro 0.16.1 and PsychoPy 2024.2.4. The stimuli were presented on an LG UltraGear 27GN950 monitor with a spatial resolution of 3,840 × 2,160 pixels and a temporal resolution of 144 Hz. We used a chin rest to fix the observers’ distance to the monitor at a distance of 55 cm. Thus, the monitor covered about 57 degrees of visual angle. All experiments took place in a darkened laboratory of the AG Perception at TU Darmstadt.

#### Analysis

For the RMSE results of the simulations (Figure 3B), the model predictions for the stimulus pool (Figure 4C), the parameter vectors (Figure 4B), and the faces in Figure 5, we computed an MCMC posterior for the psychometric function model and used the mean over the function parameters as the model for analysis. We generated faces with *σ*^*−*1^(0.0001) as very female faces and *σ*^*−*1^(0.9999) as very male faces, where *σ*^*−*1^(*x*) is the inverse of the logistic function *σ*(*x*). The entropy results in Figure 3A and Figure 4A are also for MCMC posteriors that were computed after data collection was finished.

### Sequential dependencies in adaptive vs baseline blocks

Most statistical treatments of psychometric performance assume that response sequences are independent and identically-distributed. However, empirical data often violate this assumption to some degree, with responses showing inter-trial or sequential dependencies, where the current response can be predicted from the previous responses (Fischer and Whitney, 2014). This typically causes statistical estimators that assume independence to underestimate uncertainty in the data, yielding confidence intervals that are too small (Fründ et al., 2011; Schütt et al., 2016; Fründ et al., 2014).

One reasonable concern about employing any adaptive method is that these dependencies may be stronger due to the adaptive nature of stimulus selection. Our results in the main paper demonstrate that the baseline and adaptive procedures recover similar decision functions, providing a qualitative indication that sequential effects are negligible. Here, we aim to address this question more rigorously.

To quantitatively estimate the sequential effects, we adopt an approach by Fründ et al. (2014). In essence, their approach modifies the model which is fit to the data by adding a history dependent term to the linear part of the model. One can then quantify the sequential effects as the contribution of history to the variance of the linear part. Additionally, one can compare parameter estimates obtained with and without the history term.

### Methods

The approach by Fründ et al. (2014) is limited to one dimensional psychometric functions where the label/class of a stimulus is known to the experimenters, neither of which is true for our application. The psychometric function we use is 15 dimensional (with 18 parameters) and the face stimuli do not have ground truth labels. We therefore adapt the approach from Fründ and co-authors, by adding a history dependent term to the linear part of our psychometric function:

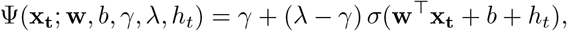

where *x*_*t*_ is the current stimulus and *h*_*t*_ captures the influence of the trial history. Depending on the value of *h*_*t*_ the predicted response is shifted towards either the male or female direction. There are multiple ways one could define *h*_*t*_. Here, we decide to include only the response given in the previous trial for reasons of simplicity, though we also tested other models and did not find qualitatively different results.

Specifically, we define:

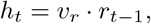

where *r*_*t −*1_ ∈ {0, 1} is the response given on the previous trial (1 = male, 0 = female), and *v*_*r*_ is a scalar coefficient fit to the data that controls the strength of the history effect. This extended model was fit separately to the data of the baseline and adaptive condition using the same MCMC procedure as the model in the main paper, resulting in a posterior distribution over all parameters including *v*_*r*_.

As a control, we fitted the extended model to a shuffled version of the experimental data, in which the order of data points was randomized. Randomization should remove sequential effects, on average.

#### Variance explained

To quantify the contribution of trial history to the model’s predictions, we computed the proportion of variance in the linear predictor **w**^*⊤*^**x**_**t**_ + *b* + *h*_*t*_ that is attributable to the history term *h*_*t*_ alone. Concretely, we compared Var(*h*_*t*_) to the total variance Var(**w**^*⊤*^**x**_*t*_ + *b* + *h*_*t*_) across all collected trials, using the posterior mean as parameter estimates.

#### Parameter shift

To assess whether ignoring trial history distorts parameter recovery, we compared the mean of the posterior distributions of the core model parameters **w**, *b* obtained with and without the history term *v*_*r*_. If sequential effects were consequential, one would expect shifts in these means when the history term is included or omitted.

### Results

The coefficient describing the influence of the previous response on the current response is consistently larger in conditions with the original sequential data than in conditions where the data are shuffled prior to model fitting (Figure 6A). There is no clear trend, however, regarding whether the coefficient is larger in the adaptive condition than in the baseline condition: for some participants (P1, P3, and P5) the estimate is larger in the adaptive condition, while for P2 and P4 the reverse holds. Notably, all estimates are positive.

**Figure 6:**
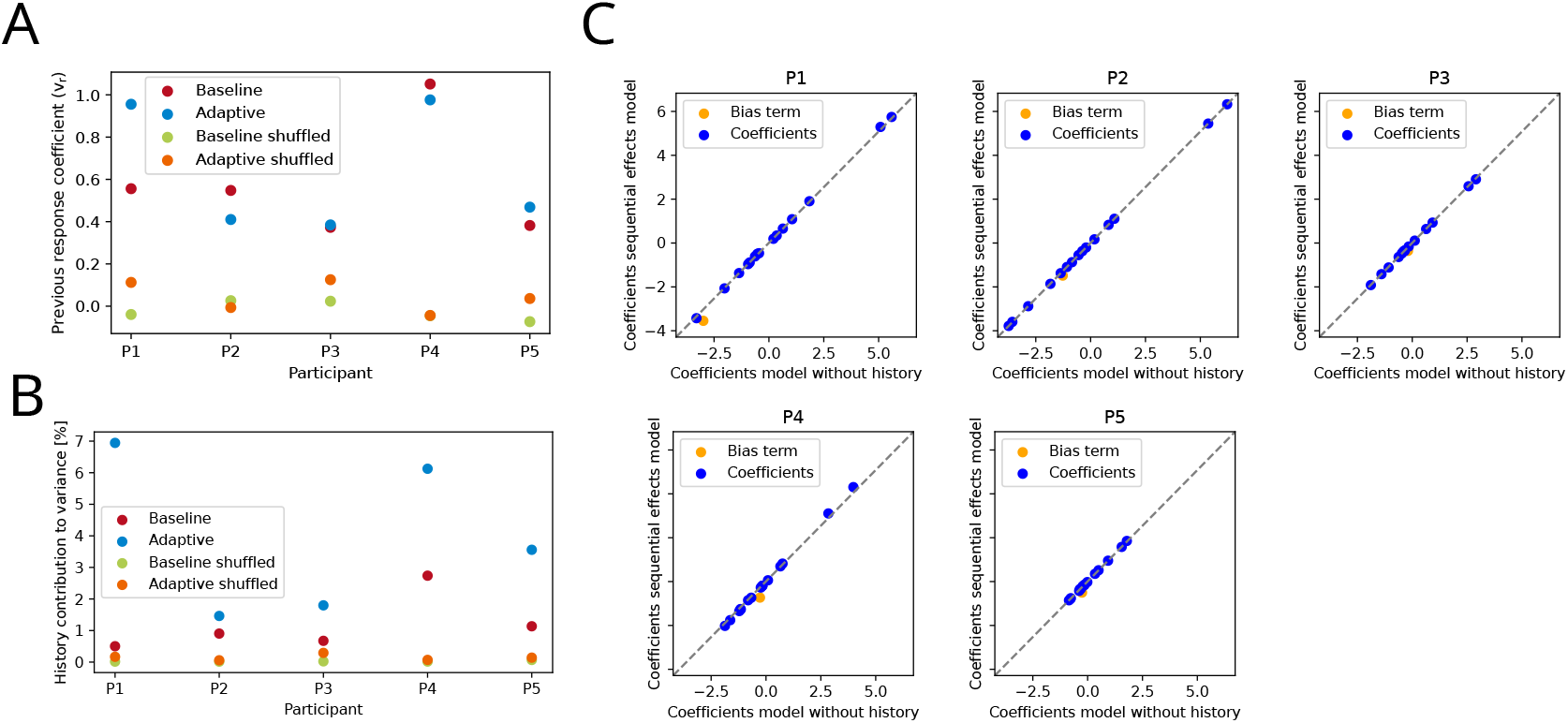
Results of sequential effects analysis. **A** Posterior means of the coefficient describing the influence of the previous response. Compared are models fitted on the baseline or adaptive condition or shuffled versions of the data of each condition. The history contribution of the models fitted on shuffled data should be close to zero. **B** Contribution of the history term to the total variance of linear part of the psychometric model. **C** Posterior means for the coefficients of the psychometric model fitted on the data collected with HOPE. We compare the coefficients of the model which includes the history term to the coefficient of the model which does not include a history term.

For both the baseline and the adaptive condition some of the variance in the linear model term can be attributed to the influence of the previous response (Figure 6A). For both baseline and adaptive stimulus selection in the original data, the history contribution to the variance is greater than for the models fitted on shuffled data, providing a check that the model is indeed capturing sequential effects. As expected, the history contribution for shuffled data is close to zero.

For all participants, the variance attributed to the previous response is larger in the adaptive condition than in the baseline condition. However, the vast majority of the variance still stems from the current trial, with variance attributed to the history never exceeding 7%.

Figure 6C shows that adding a history dependent term does not significantly influence the estimated coefficients of the psychometric function. The values of coefficients in both models align almost perfectly. Only the bias term is slightly smaller in models with history contribution.

### Discussion

The results indicate that in both our conditions (baseline and adaptive) sequential effects are present. This can be seen in the positive values of the fitted history coefficient and the fact that the history contribution to the variance is greater than for the shuffled data. However, while the estimates of the history coefficient are not interpretable in terms of effect size, the variance explained by the history term is. The fact that the history term explains only a small fraction of the variance in the linear predictor suggests that sequential effects are present but not dominant in our data. The fact that this influence is relatively small is also supported by the comparison of the fitted parameters of the psychometric function with and without the history term. If sequential effects were dominant, we would expect to see a significant shift in the recovered parameters when including or excluding the history term. However, we find that the recovered parameters are almost identical across both models with the only exception being a slightly smaller bias term in the model with history contribution. A smaller bias term is expected since the history term is designed in a way to always be added to the linear predictor (i.e. only if the previous response was “female” the history term is zero, otherwise it is positive).

The positive estimated values of the history coefficient indicate that the response given in the previous trial had an attractive effect on the response given in the current trial. In other words, if a participant responded “male” in the previous trial, they were more likely to respond “male” in the current trial, and vice versa for “female” responses. This finding is consistent with previous research documenting attractive sequential effects in psychophysical tasks (Fischer and Whitney, 2014) and the perception of faces (Liberman et al., 2014).

We find that the history contribution to the variance is larger in the adaptive condition than in the baseline condition. One possible explanation of this observation is that adaptive stimulus selection causes stronger reliance on previous responses. An alternative explanation, which we favor, is that because the adaptive procedure selects stimuli near the decision boundary more often than the baseline, the variance of the stimulus-driven component of the linear predictor is reduced, because stimuli are more ambiguous on average. When this component has lower variance, the history term accounts for a larger relative share, even if its absolute contribution remains the same. This interpretation is supported by the absence of a corresponding difference in the raw history coefficients.

### Robustness to Model Misspecification

A core assumption of HOPE is that the psychometric function follows a specific parametric form, in our case, a logistic function. In practice, the true generative process underlying observer behavior may deviate from this assumed form. We therefore tested whether HOPE retains its advantages over baseline sampling when the assumed parametric model is misspecified.

### Methods

To simulate model misspecification, we fitted two alternative psychometric functions to the Chicago Face Database (CFD) data: a Weibull function and a Gumbel function. We then sampled virtual observers, using the fits as means for priors of teh parameters plus lapse priors as in the other simulations. These observers were then used to generate responses in place of the logistic model used in the main simulations. HOPE, however, continued to assume a logistic generative process throughout adaptive sampling, introducing a deliberate mismatch between the true and assumed model. We evaluated performance against our baseline sampling method using two metrics: (1) the entropy of the posterior over psychometric function parameters, estimated by refitting the appropriate model (Weibull or Gumbel) to the collected data, and (2) the cosine similarity between the estimated and ground-truth parameters as a measure of parameter recovery accuracy.

### Results

Across simulations, the entropy of the posterior decreased more rapidly with HOPE than with the baseline for both the Weibull and Gumbel ground-truth models (Figure 7A and B). Parameter recovery, measured here as cosine similarity between estimated and ground-truth parameters (rather than RMSE as in the main simulations) likewise improved more quickly under adaptive sampling (Figure 7C and D). Cosine similarity increased faster in the early trials of adaptive compared to baseline sampling, with the advantage most pronounced up to approximately 2,000 trials, indicating that HOPE concentrates measurements in informative regions of the stimulus space even when operating under a misspecified model. By 3,000 trials, both methods reached comparable cosine similarity values.

**Figure 7:**
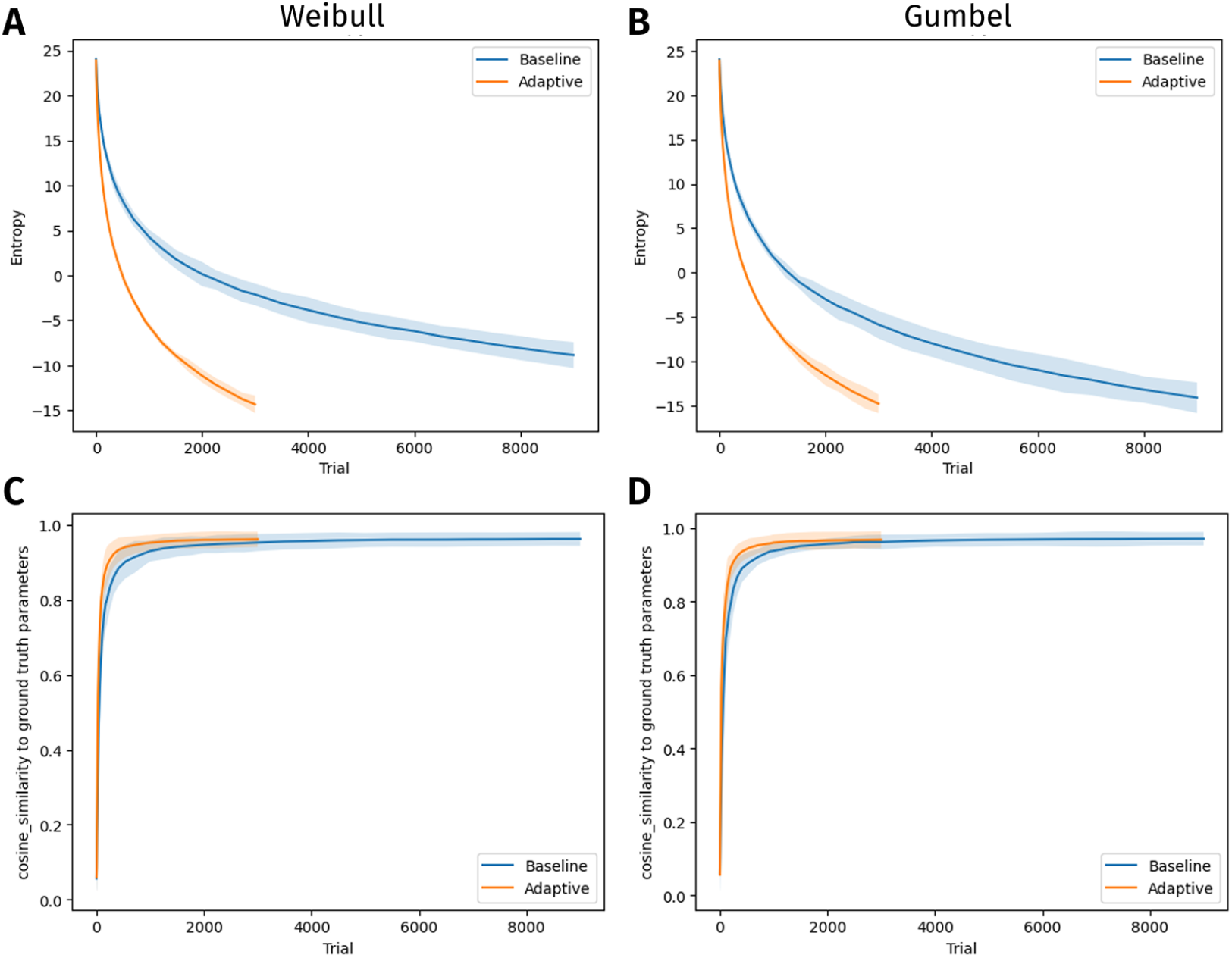
Results of model missspecification simulations. The lines represent the mean over 15 runs, respectively, the shaded areas display the standard deviation. Evaluated fits are with the underlying ground-truth functions (Weibull and Gumbel, respectively), but with HOPE assuming a logistic link function. **A** and **B**: Posterior entropy for the Weibull (**A**) and Gumbel (**B**) link functions. As in the simulations with the logistic link function, posterior entropy decreases faster for adaptive sampling. **C** and **D**: Cosine similarities between ground-truth and fitted parameters for the Weibull (**C**) and Gumbel (**D**) link functions. Cosine similarities between ground truth and fitted parameters decrease faster if stimuli are sampled adaptively.

**Figure 8:**
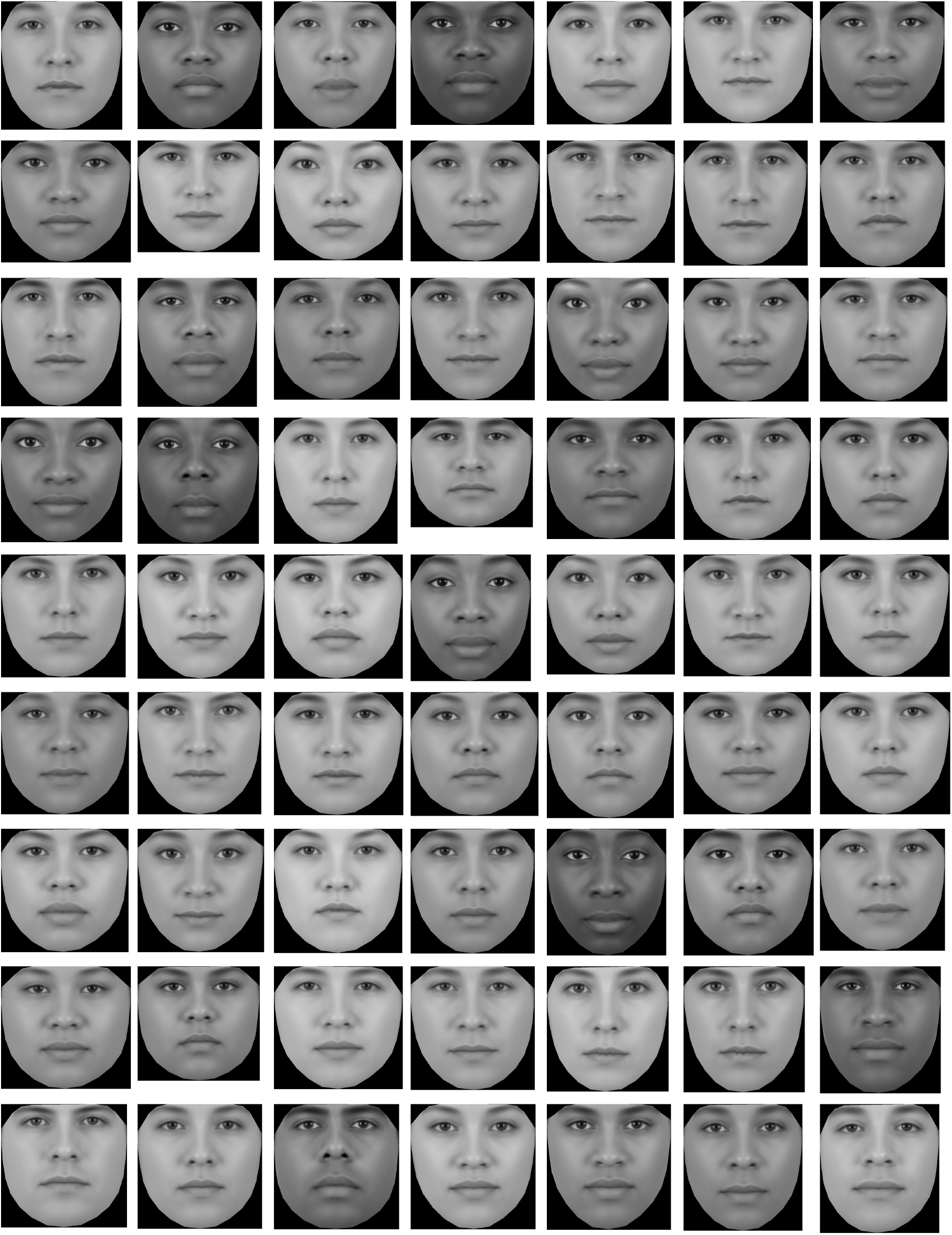
Independent examples from the stimulus pool that we used in our experiment. All faces are morphs between a random pair of one female and one male face from the Chicago Face Database (CFD) (Ma et al., 2015) and therefore do not belong to any real person. Original CFD faces are not shown.

### Discussion

These results suggest that HOPE is robust to modest misspecifications of the assumed parametric form. When the true psychometric function differs from the logistic model assumed by the adaptive method, sampling nonetheless concentrates in informative regions of the stimulus space, yielding faster posterior contraction and more efficient parameter recovery than baseline sampling. This robustness likely reflects the fact that logistic, Weibull, and Gumbel functions share similar qualitative structure: They are all sigmoidal. Therefore, the adaptive sampling strategy learned under one form transfers reasonably well to the others. Larger or more structurally distinct deviations from the assumed model may pose greater challenges, which we did not evaluate here.

### Perceived gender categorization: Example stimuli

Here we provide additional example stimuli from the perceived gender categorization experiment (Figure 8).

## Footnotes

1 Kujala and Lukka proposed pairing particle-filter-based estimation of a posterior distribution with MCMC steps to prevent particle degeneracy. They proposed this for two stimulus features (four parameter dimensions), tested it in simulation, and speculated that it should scale well to higher dimensions. It does.

